# Respiratory Muscle Training Improves Speech and Pulmonary Function in COPD Patients in a Home Health Setting - a Pilot Study

**DOI:** 10.1101/523746

**Authors:** N Bausek, L Havenga, S Aldarondo

**Affiliations:** CSO, PN Medical, Cocoa Beach FL; Asst. VP Therapy Innovation, Amedisys Home Health, Baton Rouge, LA; Med. Dir. Respiratory Care Services, Advent Health, Orlando, FL

## Abstract

- Home-based COPD managment, as provided by certified home-health providers, represents a cornerstone of patient care.
- Resporatory muscle trainig (RMT) reduces symptoms of COPD and improves underlying respiratory muscle weakness, and may be a beneficial adjunct of standard of care treatment plans.
- This 4 Week pilot study shows that RMT in combination with a standard of care home-based COPD management program can improve pulmonary and speech functions.

## Introduction

COPD is a progressive obstructive disorder, affecting more than 15 million people in the US. The cardinal symptoms of COPD are dyspnea and limited exercise tolerance, which are to a significant extent caused by respiratory muscle weakness. Approximately 19% of patients with moderate to severe COPD experience acute exacerbations, which are critical episodes of the disease requiring immediate care and hospitalization. In high-income countries, COPD is the third leading cause of death, causing 3.8% of all deaths [1] [2] [4] [3].

Treatment strategies for COPD include bronchodilators and rehabilitation to improve exercise capacity. However, the underlying respiratory muscle weakness is not addressed by current pharmacological or rehabilitative COPD management approaches.

Due to the progressive course of COPD and its burden on the health system, long-term self-management of COPD patients is critical. While patients usually experience in-hospital care during the acute stage of the disease, management of the chronic stages of COPD is often neglected, contributing to rapid progression and worsening of symptoms. Institutionalized pulmonary rehabilitation is an effective method to improve symptoms of COPD and to increase exercise capacity and QOL. However, pulmonary rehabilitation is only available to approximately 2% of eligible COPD patients [4].

Home-based exercise regimes offer a valuable alternative, and have proven to be as effective as hospital-based or outpatient pulmonary rehabilitation [5–7]. Home-based COPD management interventions such as patient activation, active monitoring of adherence, care coordination and medical management reduce sedentary behavior and increase physical activity levels. Furthermore, home-based COPD management reduces healthcare utilization, acute care hospitalization days, and mortality [8,9].

Optimal management of COPD patients includes pulmonary rehabilitation and counter measures to decrease the respiratory muscle weakness underlying the disease. Respiratory muscle training (RMT) is a drug-free therapeutic method that triggers respiratory muscle hypertrophy and improved functioning by loading the muscles during training. In resistive RMT, the air flow generated during the breathing cycle is forced through different sized apertures, adding resistance to the flow path, thereby loading the entire pressure curve of the breath flow. Intensity of RMT and workload of on the respiratory muscles is increased with decreasing aperture size [10].

This pilot study investigated the effectiveness of RMT on clinical parameters of COPD patients in a home-health setting, with the aim of exploring the potential of home-based RMT management to emulate the benefits of institutionalized pulmonary rehabilitation.

## Objectives of the Pilot Study

We evaluated data collected during a 4 week quality improvement pilot study on the effect of integrating resistive RMT into the COPD empowerment program provided by one of the largest home health care providers in the US, Amedisys Healthcare. The aims of this prospective study were:

- To evaluate the impact of using resistive RMT on pulmonary and speech function in COPD patients.
- To assess feasibility and efficacy of using resistive RMT as part of the home health COPD empowerment program, which is the current standard of care.
- To collect preliminary data on the benefits of using resistive RMT in COPD patients in the home health setting to assess justification of a larger scale clinical study.

## Study Design

Subjects with a primary diagnosis of COPD were enrolled into the standard of care COPD empowerment program including resistive RMT at the start of care with Amedisys. The intervention consisted of the standard of care COPD management provided by the home health care provider and respiratory muscle training (RMT). The COPD empowerment program consists of an introduction to diaphragmatic breathing, a warm-up section, upper limb strengthening exercises, lower limb strengthening exercises, exercises supporting cardiorespiratory fitness, a warm down, as well as patient education on COPD and its management. In addition, all patients were introduced to resistive RMT using the Breather^©^ (PN Medical, Inc.) by physiotherapists trained in the use of the device. The Breather^©^ is a resistive inspiratory and expiratory muscle trainer. RMT sessions were supervised during each therapist visit, and adherence was monitored. Frequency of visits were tailored to individual patient needs, with a minimum of one (1) to two (2) visits per week. All other RMT sessions were done by the subjects without supervision.

The resistive RMT protocol consisted of two (2) sets of 10 breaths through the device, two(2) times per day, every day, for four (4) weeks. The target training effort was set by the therapist and ranged from 40% to 60% of maximum effort. Typical pressures generated by COPD patients using the device with an effort of 50% to 70% of maximum effort range from −100 cm H2O to +43 cm H2O (manufacturer’s information). Effort level was assessed subjectively based on patient ability and flow rate. Settings were adapted appropriately by the therapist and/or the subjects to reflect the set effort target. Training sessions including training intensity and perceived effort were logged into a paper training journal by the subjects and monitored by the therapists on a weekly basis.

Significance of differences in means of clinical outcomes (MPT and PEF) was calculated using paired two-sample T-Test (critical value 5%, StatPlus). Significant outcomes are outlined by asterisks (*) in table 1.

## Study Outcomes

The outcomes of the study were peak expiratory flow (PEF) and maximum phonation time (MPT). PEF was determined by a handheld peak flow meter, and were recorded in liters per minute (lpm). MPT was measured by instructing the subjects to sit in an upright position, inhale deeply, and sustain the vowel “A” for as long as possible on the exhale. Therapists recorded MPT in seconds (s). Additional outcomes collected include compliance, settings inhale and exhale, and perceived effort.

## Results and Discussion

17 subjects were enrolled in the study, for 11 out of these, PEF and MPT were collected at the beginning and the end of the study. No other patient demographics such as age, sex, name or ethnicity were collected for the purpose of the study. Table 1 provides an overview of data collection.

**Table 1:** Pre- and post RMT outcomes.

|  | Baseline | End of Study | pValue |
| --- | --- | --- | --- |
| **Subject (N)** | 17 | 17 | N/A |
| **Setting Inhale** | 1.77 | 3.73 | N/A |
| **Setting Exhale** | 1.46 | 3.64 | N/A |
| **Effort** | 49% |  | N/A |
| **Compliance** | 84% |  | N/A |
| **MPT (s)** | 5.80 | 12.10 | <0.001* |
| **PEF (lpm)** | 190.7 | 324.5 | <0.001* |

Table 1 show outcomes collected at the beginning (Baseline) and the end of the study (Endline). Expected for subject number, all values represent means. pValue was calculated using paired t-test. Significance is indicated by asterisk (*).

## Outcome PEF

In healthy subjects, peak expiratory flow (PEF) is determined by force generated by the expiratory muscles, the time required to reach maximum alveolar pressure, lung volume, and lung elasticity and recoil. In patients with compromised lung function, PEF may be an indicator of airway obstruction and COPD disease progression [11]. Increased PEF improves cough effectiveness, sputum expectoration, pulmonary hygiene, and lowers the risk of aspiration-associated pneumonia [12]. This is clinically meaningful, as dysphagia is prevalent in COPD patients, and may contribute to exacerbations of disease [13]. In the study population presented here, mean PEF was 190.70lpm (SD: 83.89) at the beginning of the study, and 324.50lpm (SD: 114.33) at the end of the study. The significant increase in PEF over the duration of the study was 92% (p<0.001). These findings are consistent with a significant increase in expiratory muscle strength due to resistive RMT. It is reasonable to speculate here that the increase in PEF may be due to a strengthening effect of the expiratory muscles, caused by exhaling against resistance through the device. However, changes in PEF may also result from modification of associated chronic endothelial airway inflammation due to external or internal alterations, such as changes in medication or infection. Direct assessment of improvements in respiratory muscle strength by assessment of maximum inspiratory and/or expiratory pressure are therefore recommended for future larger scale studies.

**Figure 1:**
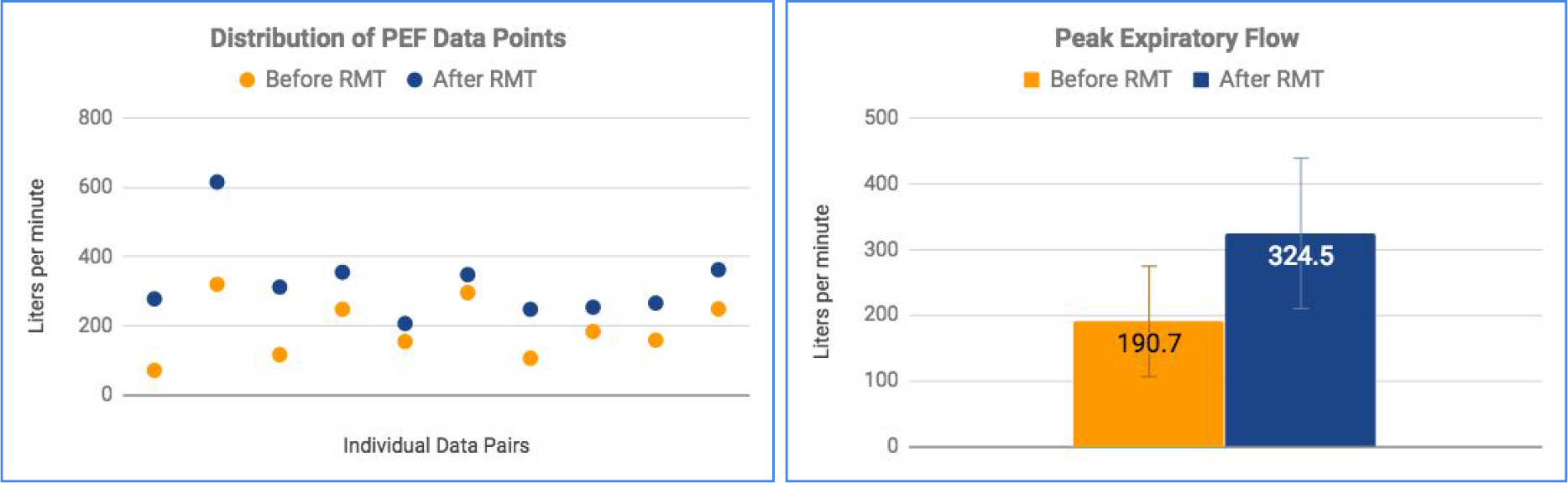
Distribution of PEF data points (left panel) show the individual collected data pairs. Orange dots indicate the values collected at the beginning of the study (pre-RMT), blue dots in direct vertical line above the orange dot indicate measurement from the same subject at the end of the study period (post-RMT), in liters per minute. Peak expiratory flow (PEF, right panel) shows average PEF pre-RMT (orange) and post-RMT (blue) values, n=11. Error bars indicate standard deviation, P-Value<0.001 (paired t-test).

## Outcome MPT

Maximum phonation time (MPT) is a simple, but highly reliable therapeutic tool to assess vocal and laryngeal function, as well as severity of dysphonia [14,15]. COPD patients have been shown to have lower MPT than age-matched controls due to higher airflow resistance and associated compromised glottal activity, affecting speech function and quality of life. Increasing MPT in COPD patients therefore has the potential to improve verbal communication quality and confidence in COPD patients [16].

Among the COPD patients investigated in the study, average MPT was 5.8 (SD: 1.62) seconds at the beginning of the study and 12.10 seconds (SD:3.18) at the end of the study. Figure 2 shows the distribution of individual data points and pairs. On average, MPT increased more than 116% over the duration of the study (p<0.001) (figure 2). Therefore, the observed increase in MPT is significant and should be confirmed in larger studies. The results indicate that RMT may increase airflow and/or glottic function, leading to improved phonation, speech breathing impacting on quality of communication. These benefits can be expected to improve social interaction, performance and ADL scores (Activity of Daily Living) of COPD patients [16].

**Figure 2:**
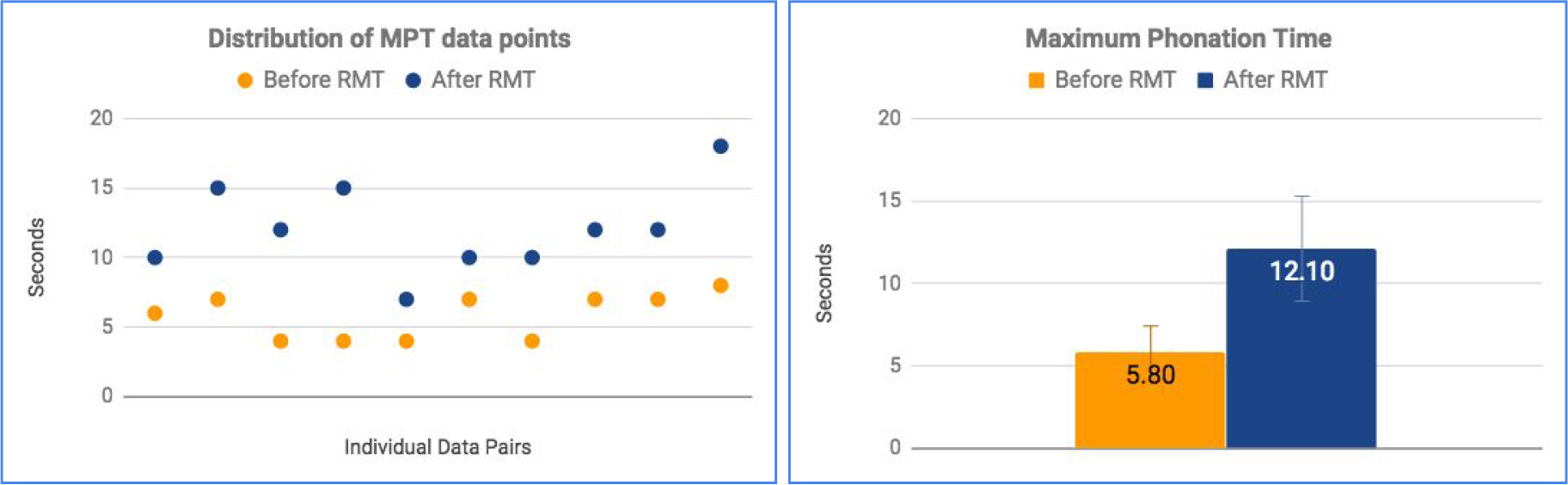
Distribution of MPT data points (left panel) show the individual collected data pairs. Orange dots indicate the values collected at the beginning of the study (pre-RMT), blue dots in direct vertical line above the orange dot indicate measurement from the same subject at the end of the study period (post-RMT), in seconds. Maximum phonation time (MPT, right panel) shows average MPT pre-RMT (orange) and post-RMT (blue), n=11. Error bars indicate standard deviation, P-Value<0.001 (paired t-test).

## Outcome Increased Respiratory Muscle Strength

Breathing against resistance increases the workload of the respiratory muscles, triggering growth of muscle fibers and increase in muscle power output. Improved respiratory muscle strength decreases the work of breathing and improves capacity to respond to higher breathing demands during exercise and ADL (Activities of Daily Living) [17]. These physiological adaptations to RMT convert to an extensive list of benefits of RMT observed across a wide range of patients, including reduced dyspnea, increased exercise tolerance and quality of life. Specific benefits include improvements of prognostic markers such as hyperinflation in COPD patients [18,19].

Figure 3 shows the increase in inhale and exhale settings over the study duration. These findings demonstrate adaptation of the respiratory muscles to RMT, consistent with an increase in diaphragm thickness and recruitment. Improved diaphragm function reduces the work of breathing and should elicit a wide range of benefits associated with increased respiratory muscle strength. In addition, increased diaphragm strength improves balance and may reduce the fall risk [20]. The increases observed over the study period are notable, with average settings reaching 3.7 for inhale and 3.6 for the exhale after 4 weeks of RMT (with both inhale and exhale settings on 1 at the start of the pilot study). As the settings range from 1 to 6 on the inhale and from 1 to 5 on the exhale, further improvements in respiratory muscle strength can be expected by continued use of RMT. Indeed, evidence from long-term studies show that benefits from RMT continue after 12 months in COPD patients [21].

**Figure 3:**
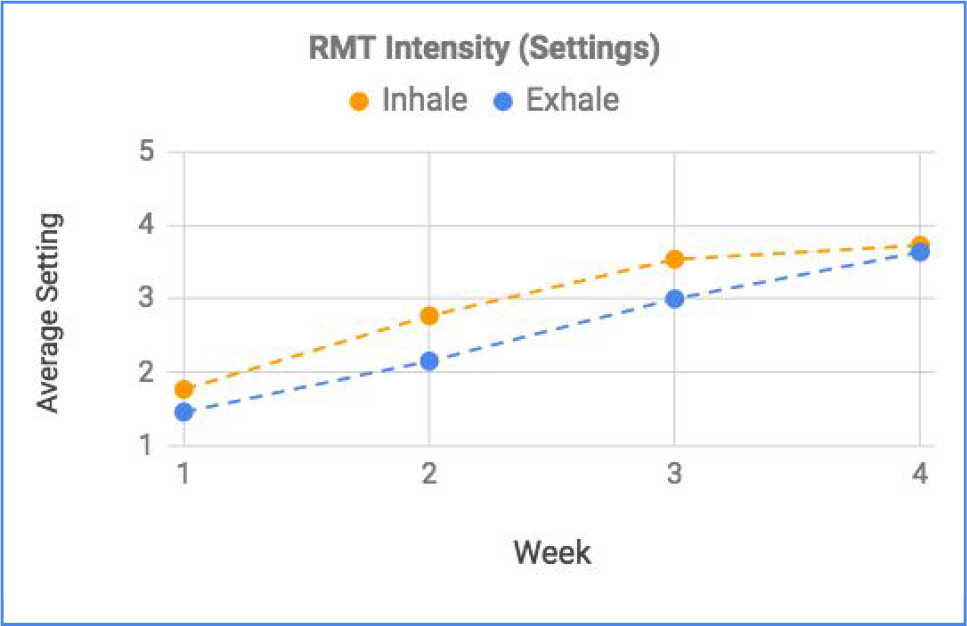
Increase of Average RMT Intensity Outlined by Increase in Settings. Orange line indicates the average increase in inhale settings (minimum: 1, maximum: 6) over the study period (4 weeks), blue line indicates the average increase in exhale settings (minimum: 1, maximum:5) over the study period.

## Outcome Compliance

12 subjects reliably reported their training sessions into the training journal. Average compliance over the study duration was 84% of sessions completed. To assess compliance, sessions that were at least 80% completed were considered. While the compliance is satisfactory and appears to yield effective results, reasons for nonadherence could be addressed to improve compliance and impact of RMT. Evidence shows that nonadherence due to low confidence in the method and benefits of RMT is the major reason for low compliance [22]. The relatively high compliance rate may be partly contributable to the use of training journals to log each session, as self-documented diaries have proven to be a useful tool for home-based rehabilitation and to provide a reliable method to monitor participation [23].

## Conclusion

This study investigates the effect of resistive RMT on the feasibility and quality of a home-based COPD management program. The results of a 4 week pilot period revealed a 116% increase in maximum phonation time and a 92% increase in peak expiratory flow. Based on these findings, the generally high adherence to the training (84% of sessions completed), and the lack of reported adverse events, the addition of resistive RMT to home-based COPD management should be considered as a feasible and beneficial option. The significant findings of this pilot study therefore justify investigation of the impact of RMT on home-based COPD management on patient outcome and readmission rates in a larger scale study.

## Limitations

The limitations of this study include the small sample size and the lack of a control group. Further limitations include the lack of direct data on respiratory muscle strength assessed by maximum inspiratory and/or expiratory pressure. For future studies, correlation of clinical data with long-term benefits such as hospitalization and readmission rates is recommended.

## Acknowledgements

The authors would like to thank Dr Tom Berlin for critical reading and insightful comments on the manuscript.

## Disclaimer

NB has received payments from PN Medical for acting as CSO. LH is an employee of Amedisys Health Care. SA has received payments from PN Medical for acting as CMO.

